# Benchmarking robust spatial transcriptomics approaches to capture the molecular landscape and pathological architecture of archived cancer tissues

**DOI:** 10.1101/2023.02.11.527941

**Authors:** Tuan Vo, Kahli Jones, Sohye Yoon, Pui Yeng Lam, Yung-Ching Kao, Chenhao Zhou, P. Prakrithi, Joanna Crawford, Shaun Walters, Ishaan Gupta, H. Peter Soyer, Kiarash Khosrotehrani, Mitchell S. Stark, Quan Nguyen

## Abstract

Applying spatial transcriptomics (ST) to explore a vast amount of formalin-fixed paraffin-embedded (FFPE) archival cancer tissues has been highly challenging due to several critical technical issues. In this work, we optimised ST protocols to generate unprecedented spatial gene expression data for FFPE skin cancer. Skin is among the most challenging tissue types for ST due to its fibrous structure and a high risk of RNAse contamination. We evaluated tissues collected from ten years to two years ago, spanning a range of tissue qualities and complexity. Technical replicates and multiple patient samples were assessed. Further, we integrated gene expression profiles with pathological information, revealing a new layer of molecular information. Such integration is powerful in cancer research and clinical applications. The data allowed us to detect the spatial expression of non-coding RNAs. Together, this work provides important technical perspectives to enable the applications of ST on archived cancer tissues.

## Introduction

As cancer is a genetically heterogeneous disease, multimodal and multiplex molecular data is increasingly being used to aid cancer diagnosis, prognosis and treatment decisions^1^. Spatial transcriptomics (ST) applications for fresh-frozen specimens has led to important findings in measuring of tumour heterogeneity^2, 3^, but this embedding type is often not suitable for clinical study sample volumes and long-term clinical follow-ups. Formalin-fixed paraffin-embedded (FFPE) tissues present a widely accessible archival biological resource and are used in all routine histopathology diagnostic laboratories^4^. Despite the many advantages of these economical, diverse and abundant samples, clinical FFPE samples are still vastly under-utilised for transcriptomic profiling due to formaldehyde cross-linking and perceived RNA degradation^5–8^.

The century-old clinical diagnostic practice based on H&E images, is qualitative and highly variable. Breakthroughs to assist pathologists to utilise the rich information in cancer biopsies are required to increase the precision of clinical decisions as well as to advance the systemic and mechanistic understanding of cancer. Unlike traditional technologies such as bulk and single-cell RNA sequencing, ST does not compromise spatial and anatomical context by tissue dissociation^9^. As a whole-tissue, spatial sequencing-based method of transcriptomic profiling, the Visium ST platform is one such technology capable of measuring ∼18,000 genes while generating histological-grade H&E images. Fluorescence In Situ Hybridisation (FISH) methods or other spatially resolved multiplex protein detection methods such as the CosMX Single Molecular Imager (SMI, NanoString), Imaging Mass Cytometry (e.g. Hyperion, Fluidigm), and Co-detection By Indexing (CODEX/PhusionCycler, Akoyabioscience) are currently available to provide single-cell spatial resolution; however, technical limitations arise when more targets are required^10–13^.

Melanoma is an aggressive heterogeneous skin cancer^14, 15^ and has been analysed by various methods, including gene expression profile^16, 17^, IHC^18^, proteomic assays^19, 20^, and fresh frozen spatial transcriptomics^21^; however, the results were still limited by the absence of histological context, low throughput and resolution, or limitations of fresh frozen tissues. In addition, skin biopsies represent the most challenging samples to obtain a consistent high-quality transcriptome^22^, especially for the old and low quality archival FFPE tissues.

In this study, we optimized both the Poly(A)-Capture Visium modified for FFPE samples (hereafter defined ‘Poly(A)-Capture protocol’) and 10X Genomics’ probe-based protocols of Visium ST (‘Probe-Capture protocol’) for human FFPE tissues from melanoma and dysplastic naevi (atypical mole). We aimed to build an FFPE ST workflow that would allow for deep interrogation of the transcriptional complexity and morphological characteristics of this challenging pathology without the need for using the limited fresh tissue samples. For the first time, we adopted, compared and combined two alternate Visium ST platforms for archived human FFPE tissue across a broad range of tissue quality and storage times. The results were also compared with the high sensitive, single-cell resolution RNAscope method. The FFPE pipeline reported here provides a high potential for revealing insights into skin cancer tissue biology.

## Materials and methods

### FFPE samples and RNA quality control

Included in this study were clinical FFPE biopsies of dysplastic neavi and melanoma, of various archival age, RNA quality and patient disease stages (Table S1). Institutional approval of experiments involving human tissues was provided by Metro South and The University of Queensland Human Research Ethics Committees HREC/17/QPAH/817, 2018000165 and 2017000318.

FFPE blocks were previously prepared in a standard procedure with fixation in 10% formalin, processed in ethanol and xylene and embedded in paraffin wax. All blocks were stored at room temperature. To assess the suitability of each sample for transcriptomic analysis, 7μm microtomed sections were collected in triplicate per sample for RNA extraction using an RNeasy FFPE Kit (#73504, Qiagen). RNA Integrity Number (RIN) and DV200 were determined by BioAnalyzer electrophoresis using an RNA 6000 Pico Kit (#5067-1513, Agilent). The DV200 metric refers to the percentage of total profiled RNA fragments greater than 200bp in length, with scores of at least 30% considered accetable for sequencing applications^23, 24^. An increasing number of fragments below this threshold in a sample is indicative of an increasing degree of RNA degradation. For this project, we selected samples with a large range of DV200 scores, with the aim of assessing the effect of FFPE RNA degradation on spatial transcriptomic data quality.

### Poly(A)-Capture

We have further optimised the protocol first developed by Gracia Villacampa, et al. ^25^, largely in terms of tissue handling and adherence, for FFPE melanoma and dysplastic naevi samples (detailed in Figure 1).

**Figure 1.**
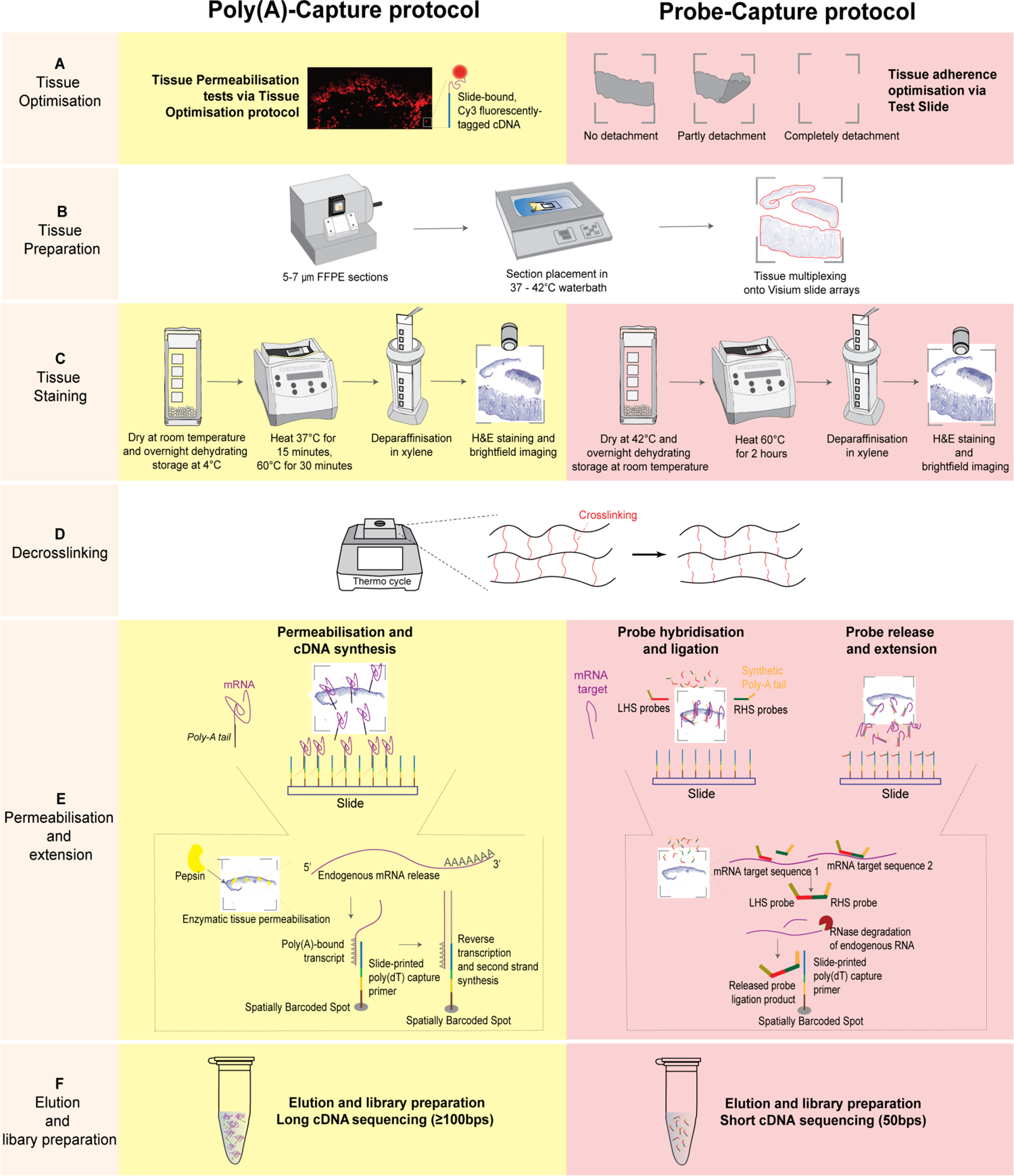
Developing and implementing protocols to perform spatial transcriptomics for FFPE tissue. (A). Poly(A)-Capture required the optimisation of tissue permeabilization step. Probe-Capture required a tissue adherence test. (B). The tissues were sectioned at 5 μm (Probe-Capture) or 7μm (Poly(A)-Capture), then floated on water bath before picking up onto the slide. The water bath was set at 37^0^C in Poly(A)-Capture protocol or at 42^0^C in Probe-Capture protocol. (C). Tissue stainning was processed in different conditions in two protocols. In Poly(A)-capture, slides were dehydrated with silica bead desiccants at room temperature for 1 hour, overnight storage at 4°C in a sealed slide-box, dried at 37°C for 15 minutes in next day, and then deparaffinised by incubated at 60°C for 30 minutes then immerse in xylene (5 minutes, twice) before H&E stanning. In Probe-capture protocol, the slides were dryied at 42°C for 3 hours, overnight stored with silica bead desiccants at room temperature, deparaffinised by incubated at 60°C for 2 hours and immersed in xylene (10 minutes, twice) before H&E stanning. (D). Decosslinking was performed in the same way (1 x TE buffer (pH 8.0) for 60 minutes at 70°C) to make RNA molecules accessible again – In poly(A)-capture, tissue was incubated in collagenase for 20 minutes at 37°C before decrosslinking. (E). In permeabilisation, the mRNA molecules or hybridized probes were released from cells and bound to the spatial oligos on the glass slide. Reverse transcription produced cDNA products in Poly(A)-Capture protocol or extended probes in Probe-Capture protocol. (F). Eluting captured molecules/probes and preparing the library for long/short cDNA sequencing. Note: RNase inhibitors were additionally included in both protocols to minimise further RNA degradation during high-temperature incubations.

#### Tissue Optimisation

The FFPE tissue sections were collected at 7μm and trimmed to include pathologist-annotated regions of interest (i.e., melanoma, stromal and lymphoid regions), and then were multiplexed per array on Visium Tissue Optimisation slides (#3000394). Slides were then dehydrated, overnight stored, then dried and deparaffinised by heat and xylene (5 minutes, twice). Tissue was then rehydrated by ethanol gradient (100% for 2 minutes, twice; 90% for 2 minutes, twice; 85% for 2 minutes). Slides were then stained with haematoxylin and eosin (H&E) and imaged using a Zeiss AxioScan Z1 slide scanner. Next, decrosslinking was performed by incubation in collagenase and then 1 x TE buffer (pH 8.0). Tissue sections were then immediately permeabilised by pepsin (0.1%) in an increasing incubation time series (5 to 40 minutes). Finally, cDNA was synthesised from the captured RNA, fluorescently labelled with cyanine 3 (Cy3), and visualised using a Leica DMi8 inverted widefield microscope.

#### Visium Spatial Gene Expression library preparation for skin cancer tissues

Following optimisation of the above conditions, FFPE blocks were sectioned and placed onto the Visium Spatial Gene Expression Slides (#2000233). Tissue was permeabilised for the duration optimised on the Tissue Optimisation slide (25 minutes). cDNA was synthesised from slide-bound poly(A) RNA *in situ*, followed by second strand synthesis and denaturation. The denatured, full-length cDNA strands were PCR amplified for 19-20 cycles. Amplified cDNA was end-repaired, A-tailed, and size-selected by SPRIselect (0.8X bead cleanup). Illumina TruSeq Read 2 sequences were ligated and standard i5 and i7 sample indexes added.

All libraries were loaded at 1.8pM onto a NextSeq500 (Illumina) and sequenced using a High Output 150 cycle kit (Illumina) at the Institute for Molecular Bioscience Sequencing Facility.

### Probe-Capture

The Probe-Capture protocol was based on the Visium Spatial Gene Expression for FFPE User Guide (CG000407, CG000408, CG000409 - 10x Genomics), with modifications as optimised for melanoma and naevus tissue.

### Tissue adherence optimisation

FFPE tissues were collected at 5μm and trimmed to include pathologist-annotated ROIs, then were multiplexed placement onto Visium Tissue Section Test Slides (#2000460). The slides were later dried, stored overnight, and deparaffinised by heat and xylene. Tissue was rehydrated by ethanol gradient following 10X protocol (CG000409), followed by H&E staining and imaging. Finally, decrosslinking was carried out in 1 x TE buffer (pH 8.0) for 1 hour at 70°C (with preconditioning in HCl).

#### Library preparation

To prepare Probe-Capture libraries for sequencing, FFPE sections were multiplexed onto Visium Spatial Gene Expression Slides. The process followed 10X user guide (CG000407, CG000409), using the whole transcriptome (18,000 protein coding genes) human probes set (#2000449, #2000450). Sequencing was performed using NovaSeq SP100.

### Multiplexed RNA in-situ hybridization with RNAscope assay

The following six target probes were designed by ACD probe design team using RNAscope Hiplex12 Reagent Kit v2 standard assay (ACD cat no. 32442): CTLA4 (ADV554341-T6), SOX10 (ADV484121-T7), Keratin8, 18 & 19 (ADV404751-T8), CD8 (ADV560391-T9), Ki67 (ADV548881-T11), CD4 (ADV605601-T12). The assay was performed according to the manufacturer’s user manual. Briefly, melanoma FFPE tissues were sectioned at 5 µm, placed on slides, and then were dried at 60°C for 2 hours before deparaffinization. Subsequently, the target retrieval step was performed followed by protein digestion with protease III. The slide was incubated with the mixture of the 6 probes or control probes for hybridisation with RNAs. After signal amplification, the slide was incubated with the RNAscope Hiplex FFPE reagent to reduce auto-fluorescence in the FFPE tissues. The signals were fluoresced and counterstained with DAPI followed by mounting with a cover slip. The imaging was performed using Zeiss LSM900 with a 63x oil objective and 5 filters (DAPI, FITC, Cy3, Cy5 and Cy7). Between imaging rounds, coverslips were removed, and fluorophores of previous imaging rounds were cleaved to enable consecutive rounds of imaging, with each round containing probes for a new set of transcripts. The single channel image at each round of image was saved and used to generate the composite image using RNAscope HiPlex Image registration Software v2.0.1.

### Data analysis

Sequencing data was mapped and demultiplexed (10x SpaceRanger), and then was analysed by a software program, stLearn^26^. The analysis consisted of: 1) processing raw data to read counts, 2) overlaying expression data with H&E tissue images, 3) performing normalisation, unsupervised clustering, 4) differential expression analysis of gene expression between spatial clusters, and 5) visualisation. We assessed heterogeneity at two levels, genes and cell types. To discriminate cell types, ST-seq derived clusters were assigned functional names by gene markers. To compare differences in cell-type composition and gene signature, we applied non-parametric tests, including Wilcoxon rank sum test and bootstrap resampling. Spatially variable genes were determined by modelling gene expression covariance with a spatial distance, implemented in the SpatialDE package.

### Noncoding RNA detection from spatial data

The data were analyzed for their long non-coding RNAs captured by the two protocols. The method described by Wang, et al. ^27^ was adopted to identify transcriptionally active regions. The pipeline uses an R package GroHMM ^28^ that utilizes a two-state hidden Markov Model to classify regions in an aligned genome as transcriptionally active or not, based on the read coverage in each bin. The position sorted BAM files generated by the 10X Spaceranger pipeline were used as inputs to the pipeline. By default, it splits the genome into non-overlapping bins of 50bp and is called transcriptionally active if reads are detected in that bin and are labelled as TARs (Transcriptionally active regions). TARs found within 500 bp apart are merged into one unit. The regions identified are then overlapped with reference gene annotations (reference annotations from 10X). The TARs overlapping with existing gene annotations are labelled aTARs (annotated TARs) and the ones falling outside gene boundaries are called uTARs (unannotated TARs). The identified novel TARs could be non-coding RNA. We overlapped these with existing databases for lncRNAs like FANTOM ^29^ and LncExpDB^30^ in a strand specific manner to identify previously reported lncRNAs.

## Results

### Optimisation of spatial transcriptomics protocols for FFPE samples

We optimised two alternate sequencing-based ST protocols for archived FFPE melanoma and dysplastic naevus tissues (Figure 1). In the Poly(A)-Capture protocol, we optimised the sectioning, deparaffinisation, decrosslinking and permeabilisation conditions. We also successfully optimised the Visium Spatial Gene Expression for FFPE tissues from 10X Genomics (‘Probe-Capture’ protocol). The use of RNA-templated ligation probes is expected to ensure high sensitivity and specificity that could be compromised for Poly(A)-Capture by relying solely on long poly(A) sequences. Figure 1 presents a step-by-step comparison between these two optimised protocols.

A primary point of optimisation commonly required for FFPE samples is that of tissue adherence to the Visium slide. Initially, we observed tissue detachment for both melanoma and naevus samples throughout deparaffinisation, staining and decrosslinking, particularly for small, overly dehydrated and fragile tissues maintaining a propensity for detachment. For the Poly(A) workflow, we performed several optimisations prior to running the Tissue Optimisation slides. Improved adherence was observed after rehydrating FFPE blocks in cold water prior to sectioning, decreasing section thickness to 7 μm, drying the slide before storing overnight with desiccator beads, and increasing the wax-melting temperature. Comparatively for the Probe-Capture workflow, a tissue adherence test replaces the tissue optimisation slide, specifically designed to minimise tissue detachment problems for experimental samples (Figure 1). For both workflows following these tests, tissue adherence was largely successful for these challenging samples.

To further optimise the Poly(A)-Capture method, we adapted the Visium Tissue Optimisation procedure for FFPE (manufacturer-designed for fresh-frozen samples) prior to library preparation (Figure S1). Exhibiting a balance between capture efficiency and lateral diffusion of RNA (decreased sharpness/specificity), we determined permeabilisation at 25 minutes to be optimal for this tissue. Optimal conditions varied between patient samples, proving Tissue Optimisation a necessity prior to library preparation for the Poly(A)-Capture protocol.

### Generating spatial transcriptomic data from Poly(A)-Capture

Following optimisations for both workflows, we performed the full sequencing library preparation on Visium Spatial Gene Expression slides. Figure 2A,B shows gene expression data from the Poly(A)-Capture workflow. By overlaying the ST data onto H&E images of the tissue taken early during the protocol, it is possible to view the number of sequencing reads and unique genes which derived from cellular/anatomical regions of interest (Figure 2A,B). From this methodology, we detected up to 2,000 genes per spot and more than 15,000 total genes per sample (Figure 2A,B), with success for both large (dysplastic naevus) and small (melanoma) tissue sections.

**Figure 2.**
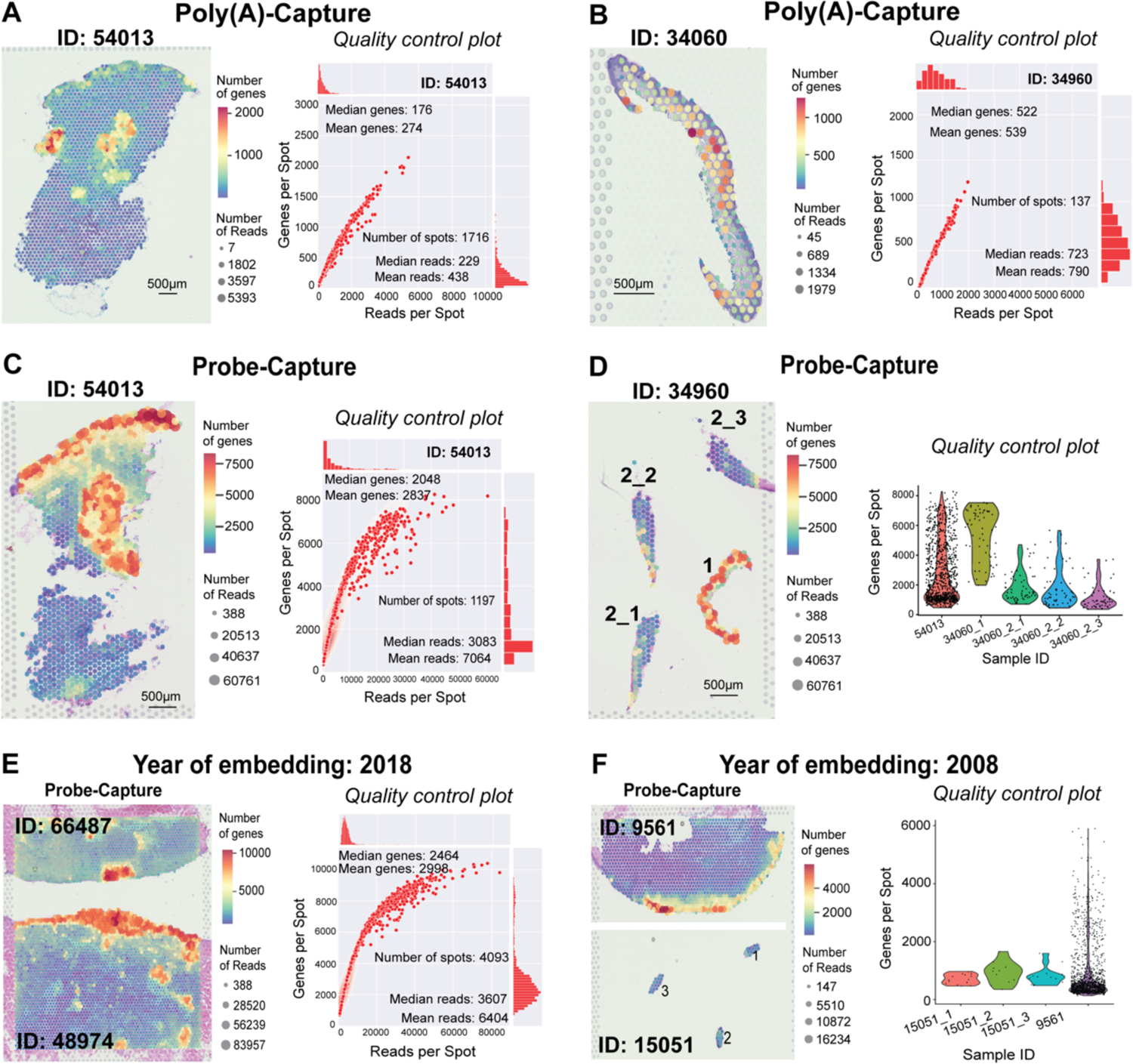
Poly(A)-Capture and Probe-Capture spatial sequencing data. (A-B). The QC for the dysplastic naevus (A) and melanoma (B) from Poly(A)-Capture protocol. (C-D). The QC for the dysplastic naevus (C) and melanoma (D) data produced by Probe-Capture method. Melanoma patient 34960 had two tissues. Tissues 34960_2 were sectioned continuously to provide triplicates on the capture area of slide (considered as technical triplicates, labelled as 34960_2_1, 34960_2_2, and 34960_2_3). (E-F). The QC for melanoma samples, which were stored for very different periods of time. The melanoma patient sample 15051 had three continuous sections from the same block, considered as three replicates which are labelled 15051_1, 15051_2, and 15051_3.

### Generating spatial transcriptomic data from Probe-Capture, assessing performance across tissue conditions and archival time

For comparison of Probe-Capture and Poly(A)-Capture protocols, we selected the same tissue blocks for analysis (i.e., adjacent sections, patient 54013 dysplastic naevus and 34960 melanoma). As expected, we observed a marked increase in the number of genes detected per spot (Figure 2). For the sample replicates across each protocol, we could detect on average 2,837 genes per spot, with up to 8,000 genes per spot using Probe-Capture (Figure 2). We also assessed technical accuracy of the method and intra-patient variation by analysing three technical replicates (adjacent sections of the same tissue piece, 34960_2_1/2/3) and two different biopsies from the same patient (34960_1 vs 34960_2) (Figure 2C,D). Capture results were consistent across technical replicates, demonstrated by the similar number of genes per spot, much more similar compared to that in other tissue sections, even for those from the same patient (Figure 2D). As expected, there was a clear disparity in the number of genes detected per spot between different biopsies of the same patient (Figure 2D), indicating that selection of biopsies with variable morphology and anatomical details, even when derived from the same clinical sample, can impact efficiency of ST.

A challenging aspect of translational research, particularly for retrospective studies, is analysing clinical samples of variable storage times, storage conditions and processing methods, any of which can negatively impact RNA quality. In this project, we assessed the efficiency of the ST methods to analyse clinical samples collected 4-14 years prior. Newer tissues (66487 and 48974, from 2018) had average DV200 scores of 70%, while older samples (9561 and 15051, from 2008) had average scores of only 31%, clearly demonstrating an impact of FFPE sample age on RNA quality. Using Probe-Capture Visium, we detected substantially more genes in the newer samples, with up to 10,000 genes per spot (Figure 2E). In contrast, the older (and more degraded) samples yielded a maximum of 6,000 genes per spot (Figure 2E). As anticiptaed, the data shows that samples of lower initial RNA quality indeed yielded decreased unique gene counts – a major consideration moving forward with FFPE ST. Of note, despite the reduction in the gene detection sensitivity, the information from these samples was sufficient for mapping cells consistently to histological annotation. Additionally, similar to the replicates shown in Figure 2D, we again saw consistency in QC between three adjacent replicate sections of the 15051 patient (Figure 2F). This suggests that the data from spatial profiling was reproducible.

### Detecting noncoding RNA from Poly(A)-Capture and Probe-Captured Data

While most of the analyses for spatial transcriptomics data have been focusing on protein coding genes, there is a huge potential to detect long non-coding RNAs (lncRNA) in the tissue. Successful detection on lncRNA spatially will allow to associate their spatial expression patterns with morphological features. Analysing multiple replicates, we found that the polyA-capture protocol detected a large number of lncRNA (>9000 lncRNA per sample), much higher than those detected by the probe-capture protocol (Figure 3). Importantly, more than 50% of the detected lncRNA also present in the two well-curated lncRNA databases, LncExpDB and FANTOM, suggesting that the lncRNAs in the spatial datasets are likely true lncRNA. Thus, the polyA-capture protocol, although detected fewer genes in total, can find a significant number of lncRNAs. Overall, this suggests the complementarity between the two protocols and that the poly(A)-capture protocol can have important roles that the probe-capture protocol alone could not meet.

**Figure 3.**
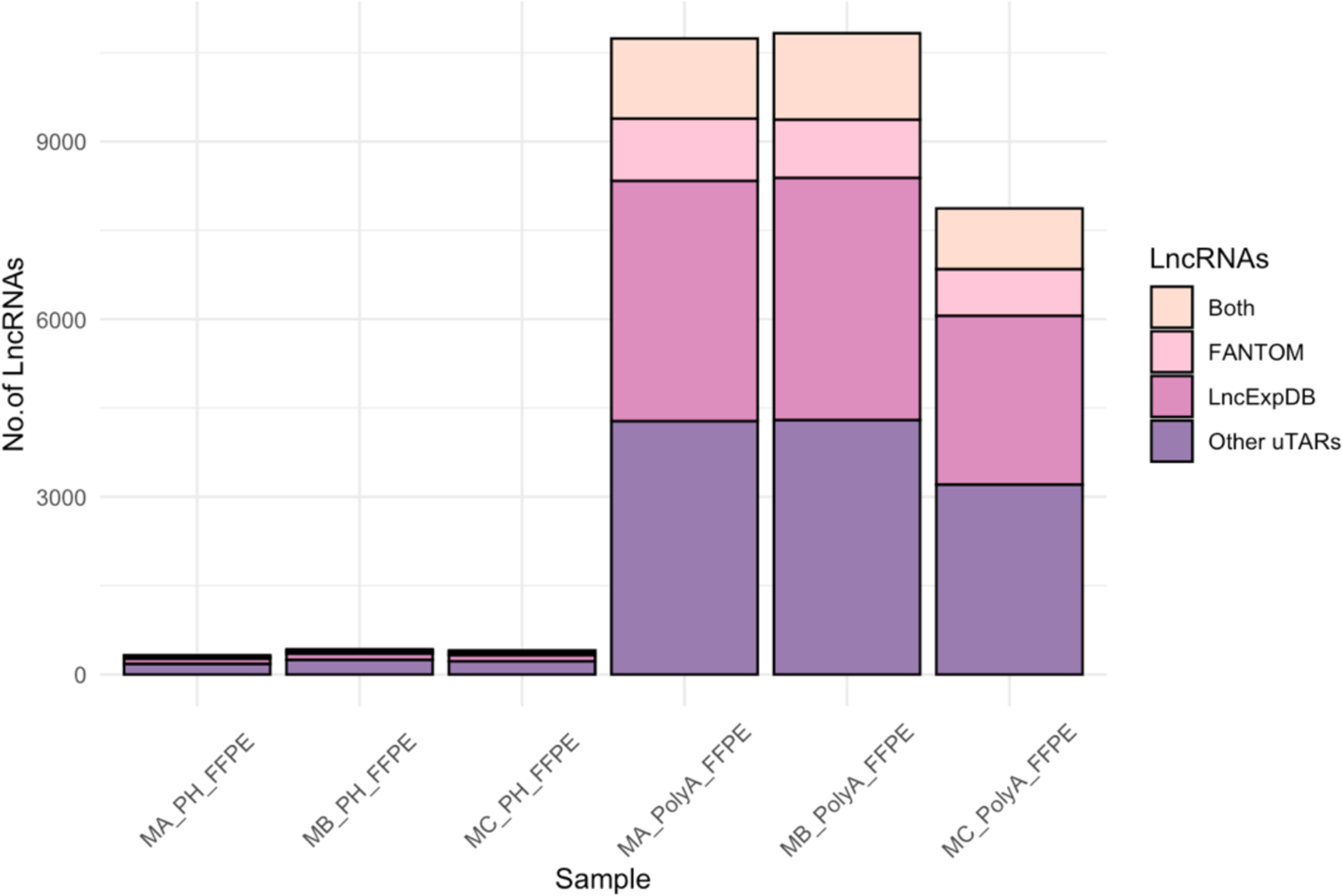
Detection of noncoding RNA (lncRNA) in melanoma samples by the polyA-capture (polyA_FFPE) and probe-capture (PH_FFPE) protocols. The captured lncRNAs are classified as belonging to previously reported lncRNA in the lncExpDB and/or FANTOM protocol. Replicates are shown as MA, MB, and MC representing samples from three melanoma patients.

### Characterising heterogeneity within the FFPE tissues

The spatial transcriptomics data of the FFPE samples that were 4 years to 14 years of storage both could accurately map cell types to the tissue. Here we assessed two skin disease stages, a dysplastic naevus and melanoma. Of note, three technical replicates as consecutive sections from the same block were included to assess technical variation and reproducibility. For the dysplastic naevus, the unsupervised clustering shows that the data from probe-capture could lead to a higher-resolution classification of tissue types. As the manual annotation from the pathologist identified the heterogeneity of dysplastic naevus skin (Figure 4A,F; Figure S2A,B), we ran spatial clustering at spot level (one spot contains 1-9 cells). We identified four clusters in Poly(A)-Capture data and nine clusters in Probe-Capture data that overall match the manually annotated tissue types. In Poly(A)-Capture data, we defined collagen (with markers *COL1A1, COL1A2, DCN*), Sebasceous gland (*FADS2, MGST1*), Eccrine ducts (*DCD, SCFB1D2*), Keratinocytes and melanocytes (*KRT10, KRT1, TYRP1*) (Figure 4B,C). In Probe-Capture data, we detected more Lymphocytes (cluster 5 - *ACTB, TMSB4X, PNRC1*) (Figure 4G,H) within the of sebaceous gland clusters and eccrine ducts clusters. Of note, in the Poly(A)-Capture data, by sub-clustering cluster 2 (Keratinocytes and melanocytes), we could find lymphocytes (*CD74, HLA-DRB1, HLA-DRA*) and melanocytes (*PMEL, DCT, TYRP1*), (Figure 4D,E).

**Figure 4.**
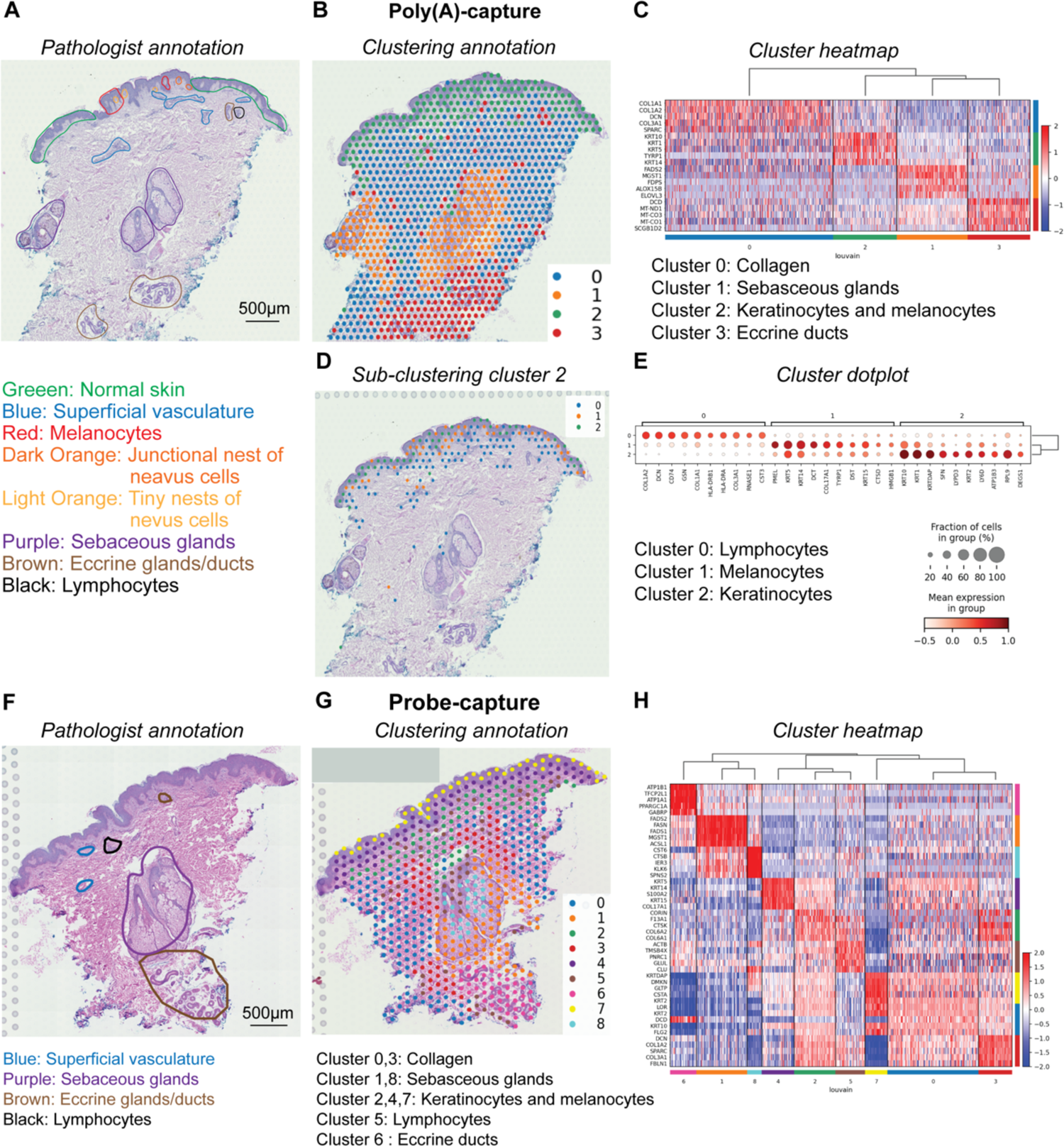
The data-driven map of heterogeneous populations on dysplastic naevus tissues. (A). The annotation of dysplastic naevus from poly(A)-capture by pathologist. Pathological annotation is shown as colour circles. (B-C). The spatial clustering (B) and heatmap (C) of dysplastic naevus from poly(A)-capture revealed the molecularly defined clusters that are heterogeneous and consistent with pathological annotation. (D-E). Spatial sub-clustering (D) and heatmap (E) of cluster 2 defined in the first round clustering of dysplastic naevus (as shown in B-C). (F). The annotation of dysplastic naevus from probe-capture by pathologist. Pathological annotation is shown as colour circles. (G-H). The clustering of dysplastic naevus tissue from probe-capture (G) shows more heterogeneity details. Heatmap (H) shows top gene markers for each cluster.

For the melanoma samples (Figure 5), the data for patient ID-48974, which was collected in 2018, contains six main clusters. Gene markers for these clusters, as shown in the heatmap, suggest cell type annotation consistent with tissue regions determined by the pathologist (Figure 5A-C, Figure S3A). We defined Melanoma (*PMEL, MLANA*), Immune infiltrates (*TRBC2, TRAC, TMSB4X*), Melanophages (*CD74, LYZ*), Keratinocytes (*KRT14, TRIM29*), Blood vessel (*<u>CAVIN1, PECAM</u>*), Collagen (*DCN, COL1A2, FBLN1*). Depending on tissue sizes and complexity, the number of clusters changed. A smaller tissue from patient ID-9561, collected in 2008, had four clusters, including Melanoma (*PMEL, TYRP1*), Immune cells (*TMSB4X, IL32*), Keratinocytes (*KRT10, KRT1, DSG1*), Collagen (*COL1A2, COL1A1, DCN*) (Figure 5D-F, Figure S3B). For the smallest tissue from patient ID-15051, with three biological replicates, there were two specific clusters consistently defined across the replicates. These two clusters are keratinocytes and melanocytes (cluster 0 – *S100A2, SPARC, TYR, MYO10*) vs epidermis (cluster 1 – *LCE2C, FLG*), consistent to the pathological annotation (Figure 5G-I, Figure S2C).

**Figure 5.**
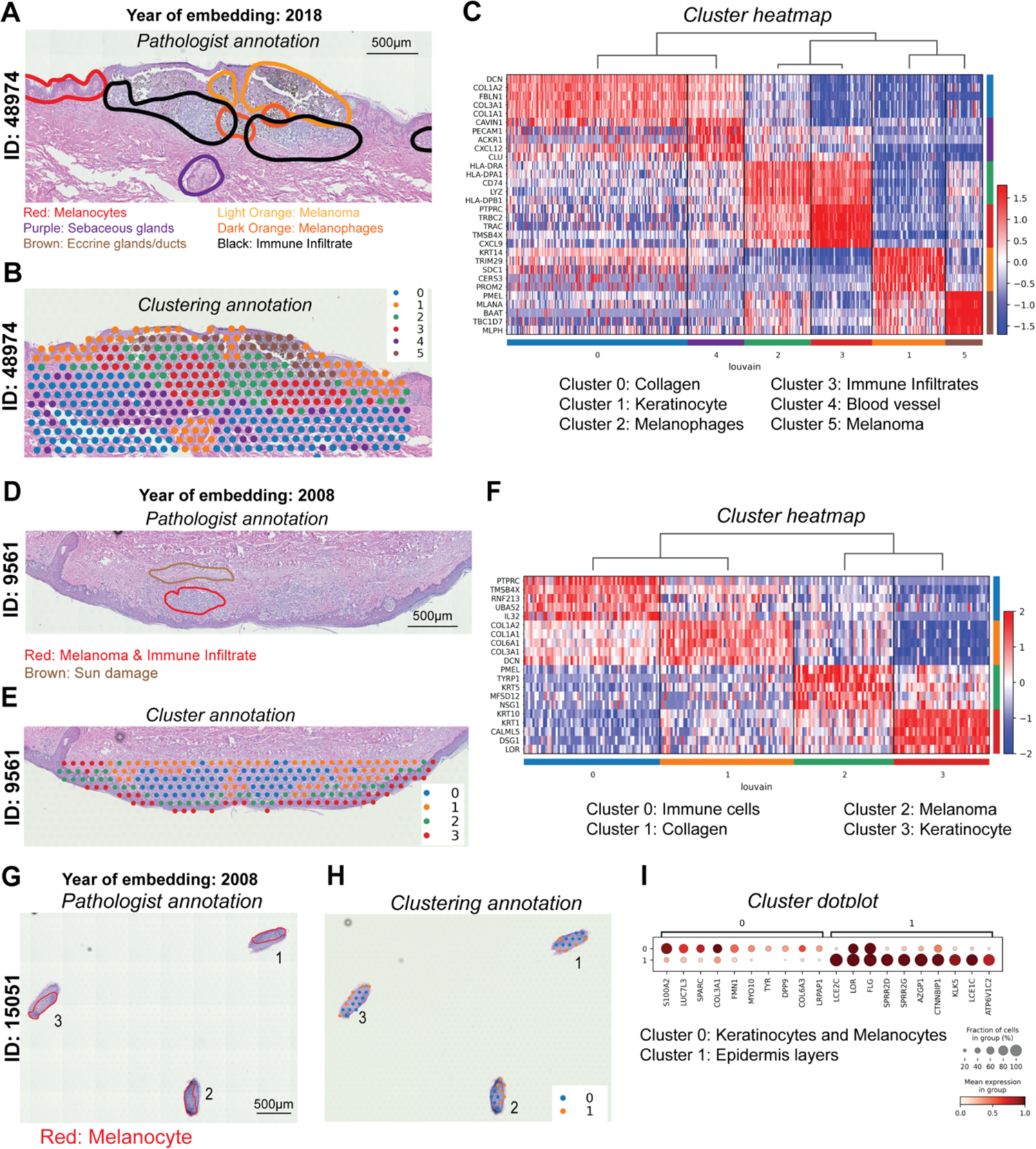
Visium Probe-Capture for melanoma FFPE samples stored at different periods of time. (A, D, and G). The pathological annotation as colour circles. (B, E and H). Corresponding clustering results from tissues in A, D, and G, respectively. (C, F and I). Heatmaps of gene marker expression for each cluster in B, E and H, respectively.

Having established the experimental protocols to robustly perform spatial transcriptomics on FFPE tissue, we next aimed to study (pre)melanoma tissue heterogeneity at gene and cell level. Based on the expression profiles of over 15,000 genes across the whole tissue section (up to 5000 spots per tissue), we identified 10 molecularly distinct cell types or functional regions for the dysplastic naevus sample (Figure S4). These cell types and regions showed spatially specific expression of gene markers, for example the pigment-cell (melanocyte) specific Premelanosome gene (PMEL) encoding melanocyte-specific type I transmembrane protein. Visual inspection of PMEL gene expression also suggested that PMEL was expressed in naevus region (Figure 2). Less known marker genes, specific to a cell type or a functional region, like the PRDX2 can be detected (Figure S5). Together, our data showed strong evidence that the spatial gene expression was able to capture tissue heterogeneity at a high resolution, across the whole tissue section and in an automated and unbiased way.

Moreover, to evaluate our findings from the FFPE ST study, we performed RNAscope assay which produced single cell resolution and high sensitivity in gene detection (Figure 6). Since the current RNAscope technology using FFPE sample is able to detect a small set of genes (up to 12 molecules), we selected six genes as markers of cancer cells and immune cells. Similar to ST experiment, we also provided the pathological annotation based on nuclei shapes and distribution from the same slide, defining immune infiltration and superficial melanomas regions (Figure 6A). Each punctate dot signal on a cell represents a single molecule in the assay. As a result, the assay established the abundant expression of SOX10 in the superficial melanoma region along with its co-expression with MKI67 (Figure 6A1). Also, the distinct co-expression of CD4 and CTLA4 was seen in the immune cell infiltrate area with a low expression of CD8 (Figure 6A2). Compared to pathological annotation, it appears that both ST and RNAscope can define the cancer and immune regions, but with much more infomration on molecular expression profiles that mark individual cell types and activities. While RNAscope provides single cell resolutions and high detection sensitivity, the ST generated data for thousands of times more genes.

**Figure 6.**
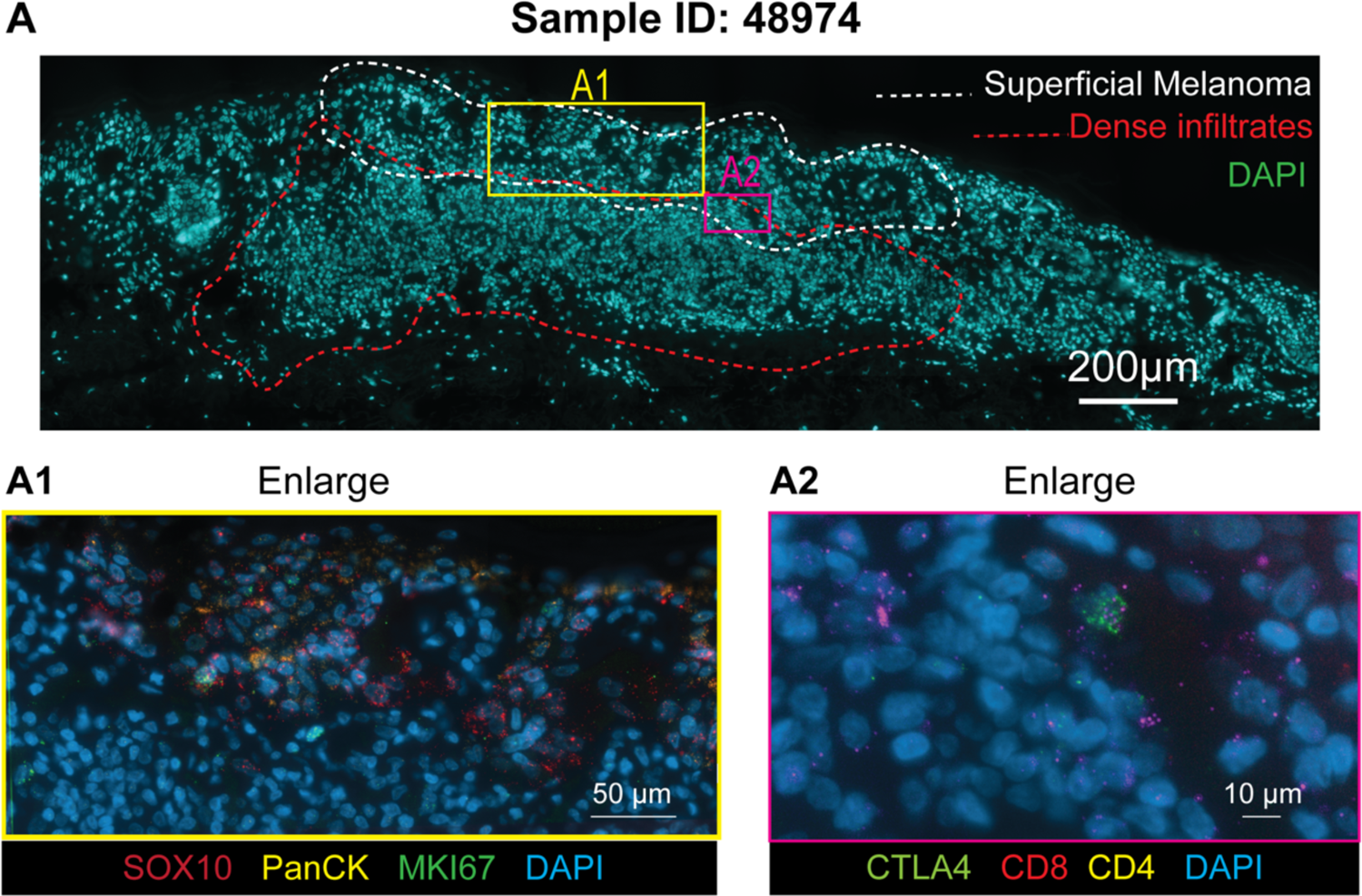
Targeted RNA molecule expression at a single cell level using RNAscope assay. (A). An overview of the section with the nuclei stained with DAPI and pathological annotation circled by white and red lines. The zoomed-in of a superficial melanoma region, showing two windows A1 and A2. (A1). With the display of cancer markers SOX10, PanCK, MKI67. These genes are expressed in the melanoma metastasis region near the epithelial layers. Each punctate dot represents a single copy of an mRNA molecule. (A2). The expression of CD4 T cell marker (CD4, CTL4A) and CD8 T cell is observed in the immune cell infiltration area.

## Discussion

Archived FFPE tissue samples, a worldwide standard in pathology departments, provides an invaluable resource for molecular research due to enormous number of biobanked collections^31–35^. Despite the vast potential for pathological applications, ST has not been popular for these samples due to nucleic acid crosslinking, molecular degradation, and tissue-slide detachment^8, 35^. In this study, we established two alternate ST methods to overcome these challenges with FFPE tissues. Importantly, we assessed tissues of variable sizes, archival times, cancer progression level and RNA quality across biological and technical replicates.

In clinical practice, manual observation of FFPE melanoma tissues by pathologists is often limited to assess tumour heterogeneity, in turn meaning that accurate diagnosis and effective treatment plans can be obstructed^36–41^. Current common spatial techniques^42–46^ can on average detect less than 100 proteins and fewer than 300 gene markers. Comparatively, Visium is an ST technology that is capable of measuring the spatial whole transcriptome and near single-cell resolution^47–49^ and at the same time generating histological-grade tissue images. We optimised the Poly(A)-Capture protocol as this method can capture RNA that are not in a predefined probeset, thereby providing missing information like the expression of lncRNA or in the case of detecting RNA from a species without predesigned probes. The gene detection capacity of the two FFPE protocols reported here can be thousands of times higher than classic pathology techniques. The Probe-Capture protocol detected more genes with increased sensitivity, but missed genes not in the panel, especially lncRNA. This is important because, lncRNAs plays an important role in melanoma development including proliferation, invasion, and apoptosis^50^. Our protocols worked with challenging FFPE skin tissues older than 12 years old, with high degradation (DV200 <30%) (Figures1, 5, 6). We have tested numerous sectioning and storage conditions, as well as section thickness to improve section adhesion^8^, balancing the improved adherence and protection of RNA quality. Moreover, since cost is a major barrier to applying ST, we also validated the option to multiplex tissue samples into Visium capture arrays for space maximisation. In this way, we were able to analyse up to nine tissue sections per slide, rather than a standard four.

From the thorough assessment of these protocols, we suggested that for discovery purposes, an unbiased approach FFPE poly(A)-capture approach should be applied as it detect all genes, including lncRNA. By comparing multiple replicates, we found that both protocols have high reproducibility, with much less technical variation compared to biological differences. Thus to capture cancer heterogeneity we recommend that biological replicates are more important than technical replicates. We also demonstrated a multiplexing strategy to practically reduce cost and thus allowing to increase sample size. For low throughput confirmation of the result, we suggest using RNAscope with high sensitivity and resolution. These comprehensive results to provide new approaches to processing old and degraded FFPE tissues for spatial transcriptomics open a new horizon to explore skin cancer tissue biology.

## Acknowledgments

We gratefully acknowledge the contribution of IMB sequencing facility - Institute for Molecular Bioscience. This work was partly funded by Genome Innovation Hub (GIH) at the University of Queensland.

## Author contributions

QN, MSS, KK conceived the study. TV, KJ, SY, PYL, JC performed experiments with the help of SW. QN, TV, PP, IG, performed data analysis. YCK, CZ, KK, MSS provided samples and ethic management. PS annotated H&E tissue sections. QN, TV, KJ, SY, MSS wrote the manuscipts. All authors have read and approved the manuscript.

## Data and code availability

Datasets supporting this manuscript are available at Zenodo, DOI:10.5281/zenodo.7475873 and code supporting this manuscript used our stLearn spatial transcriptomics analysis software available at https://stlearn.readthedocs.io/en/latest/.

---

**Figure S1.**
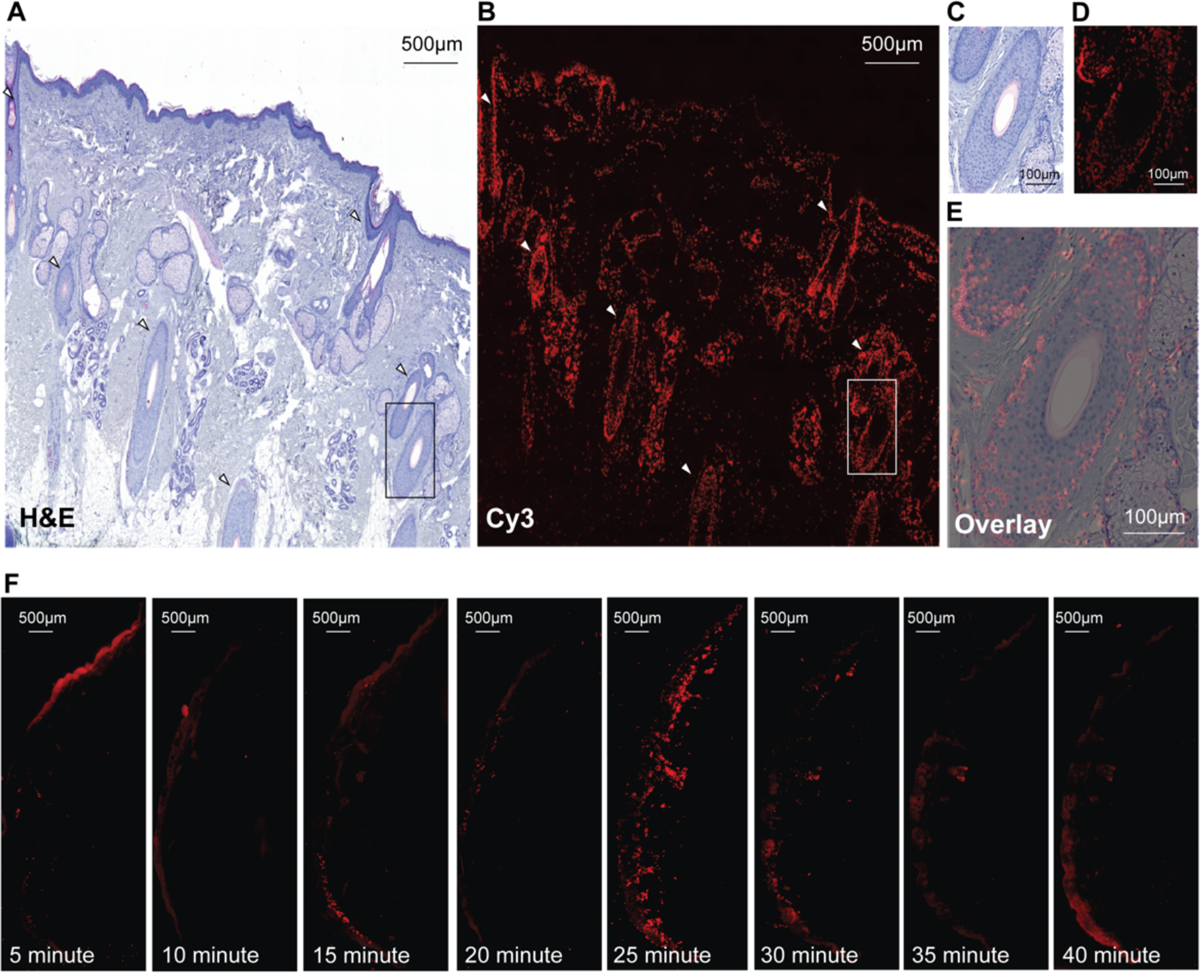
Tissue Optimisation experiment performed prior to the Poly(A)-Capture workflow. (A). Brightfield (H&E-stained) image of the tissue section. (B). Fluorescent (Cy3-tagged, poly(dT)-bound cDNA) image of the issue section. (C). Box denotes an enlarged region on the brightfield image. (D). Box denotes an enlarged region on the fluorescent image. (E). Overlays can be used as a measure of quality control by assessing that Cy3 signal is consistent to H&E morphology, with cDNA concentrated to the densely nucleated follicular tissue. (F). Cy3 images as a time series of tissue section permeabilisations, beginning with 5 minutes and proceeding to 40 minutes (incubation with 0.1% pepsin). The 25 minute permeabilisation was chosen as optimal from the series, with highly concentrated poly(dT)-bound cDNA evidenced as the most intense and tissue-specific Cy3 signal.

**Figure S2.**
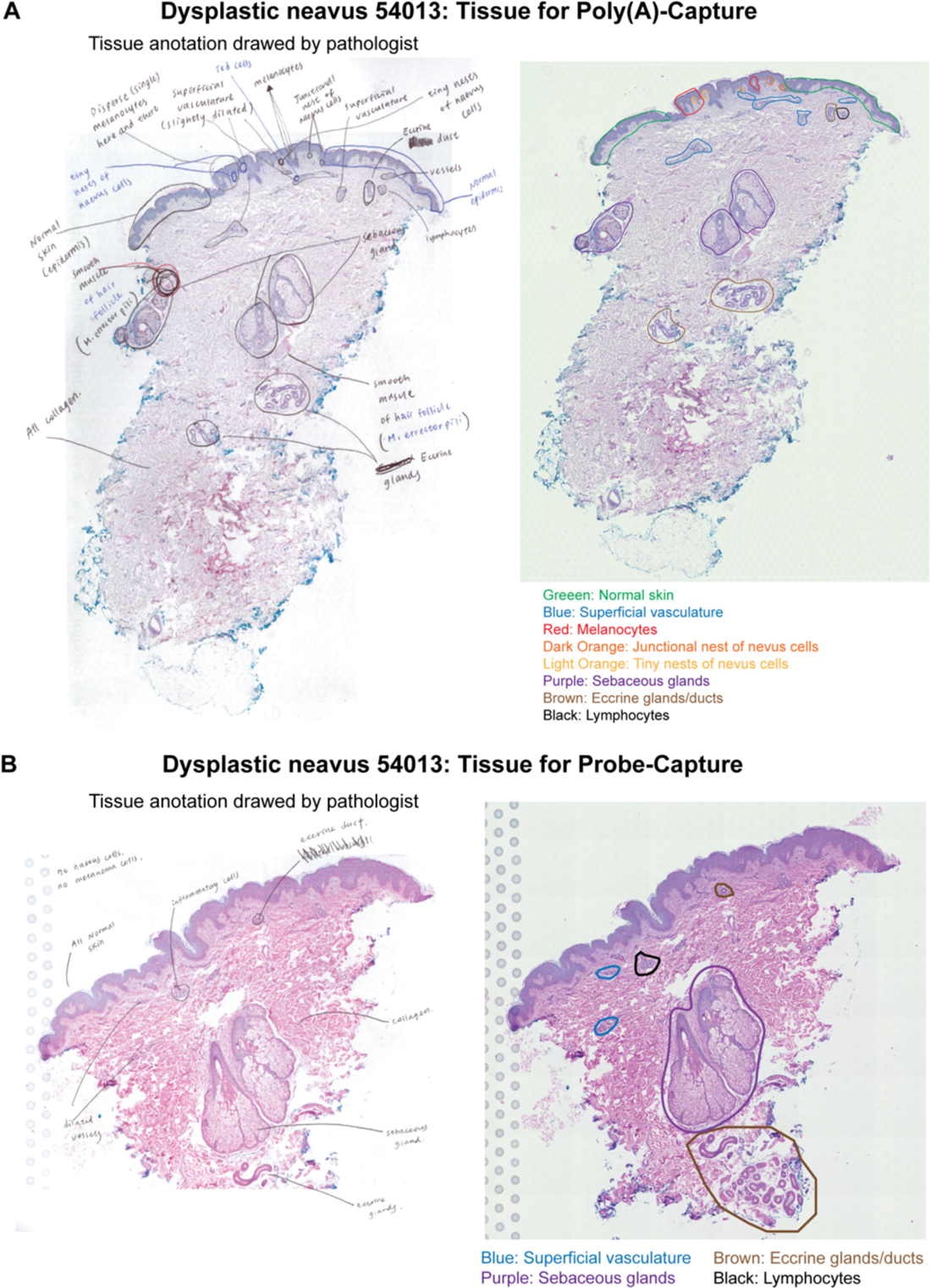
The pathological annotational of the Dysplastic naevus section used in Poly(A)-Capture protocol and Probe-Capture protocol. (A). Dysplastic naevus section used for poly(A)-capture protocol. Left is the original annotation and right is the transfer of the selected regions with colour coding. **(B)**. Dysplastic naevus section from the same block, but was cut deeper, used for probe-capture protocol. The annotation from left is transferred to the right with colour codes.

**Figure S3.**
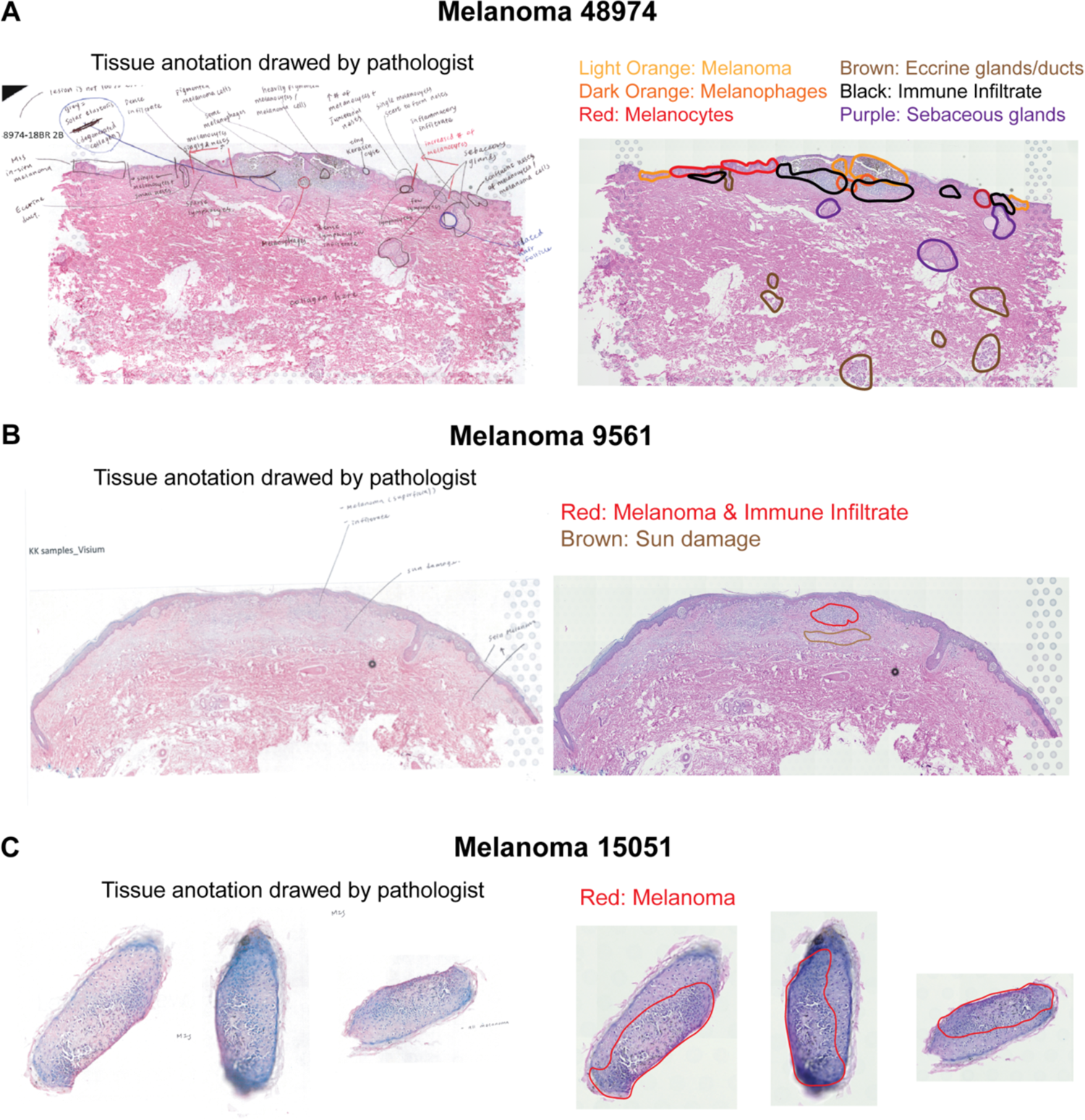
The pathological annotational of the melanoma tissue sections that used in this paper. (A) Annotation for patient 48974. The six regions are coloured coded and transferred from left to right. (B) Annotation for patient 9561. The annotated melanoma and sun-damaged regions are transferred from left to right images. (C) Annotation for patient 15051. Three sections are three technical replicates.

**Figure S4.**
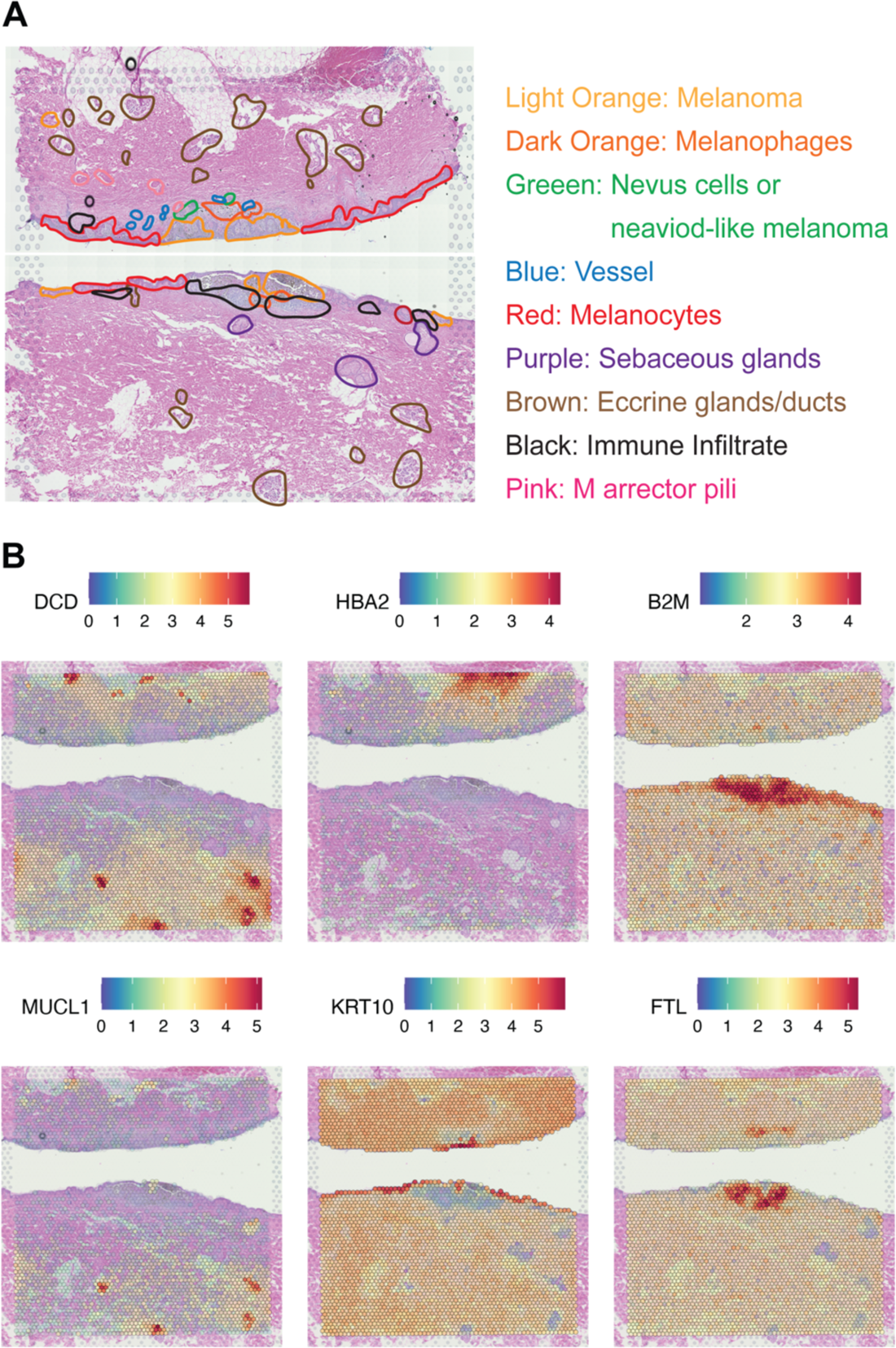
Spatial heterogeneity at gene level. (A). Pathological annotation for the two tissues. (B). The heatmap gradient colours show the expression level across the tissue section. The top six most spatially variable genes are shown. These genes were identified without human inputs from prior knowledge.

**Figure S5.**
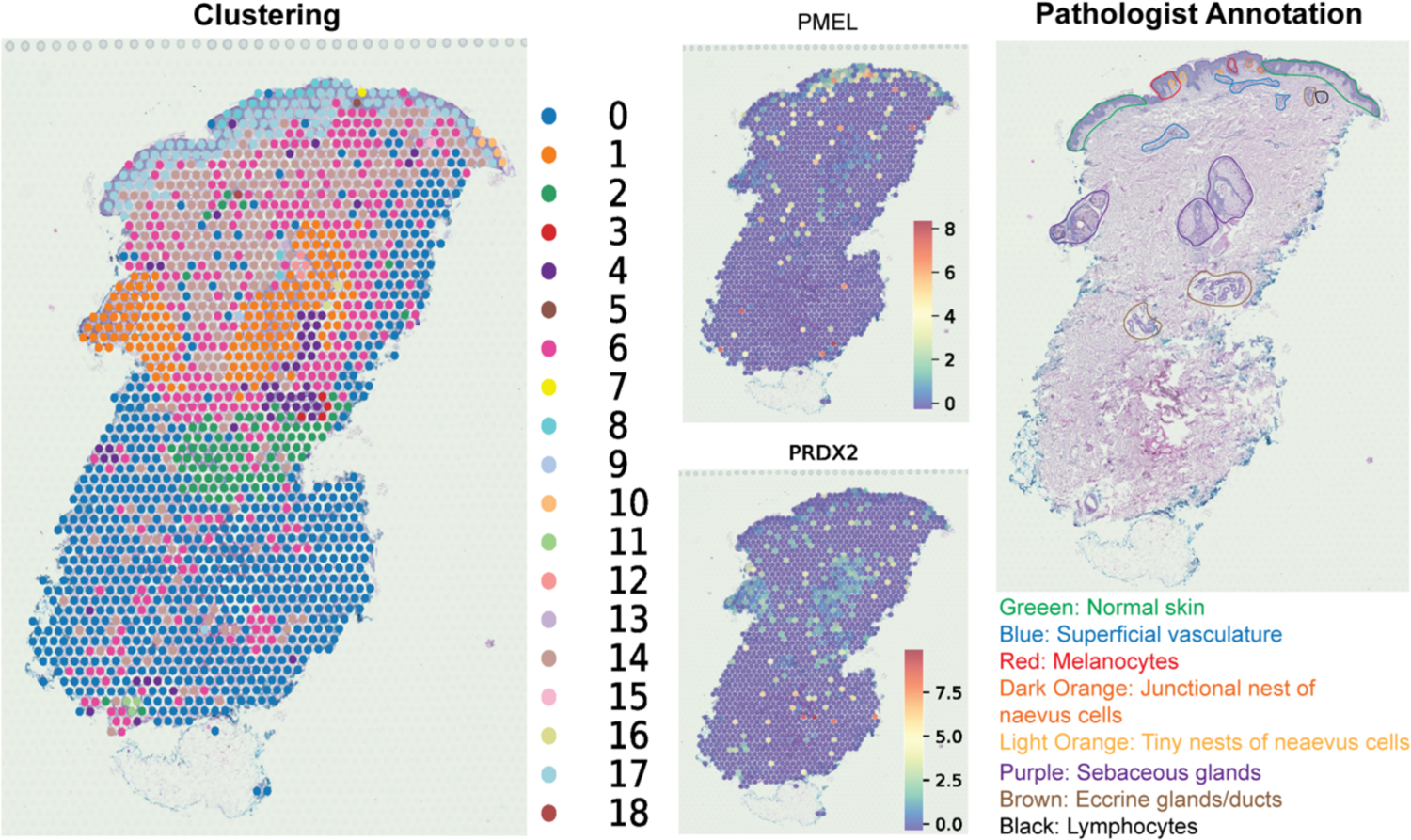
Spatial heterogeneity at gene level. The clustering results are shown on the left, histopathological on the right. The heatmap gradient colours in the middle show the expression level of two melanoma markers across the tissue section.

**Table S1.** Information of FFPE tissue samples used in this study

